# Slow-cycling cells in glioblastoma: a specific population in the cellular mosaic of cancer stem cells

**DOI:** 10.1101/2022.01.25.477703

**Authors:** C Yang, G Tian, M Dajac, A Doty, S Wang, JH Lee, M Rahman, J Huang, BA Reynolds, MR Sarkisian, D Mitchell, LP Deleyrolle

## Abstract

Glioblastoma (GBM) exhibits populations of cells that drive tumorigenesis, treatment resistance, and disease progression. Cells with such properties have been described to express specific surface and intracellular markers or exhibit specific functional features including being slow-cycling or quiescent with the ability to generate proliferative progenies. In GBM, each of these cellular fractions was shown to harbor cardinal features of cancer stem cells ^1-7^. In this study we focus on the comparison of these cells and present evidence of great phenotypic and functional heterogeneity in brain cancer cell populations with stemness properties, especially between slow-cycling cells and cells phenotypically defined based on the expression of markers commonly used to enrich for cancer stem cells (CSCs). Our data support the cancer stem cell mosaicism model with slow-cycling cells representing critical tesserae.

## Introduction

The cancer stem cell paradigm originated from studies of acute myeloid leukemia, which contains a subpopulation of cells showing stem-like cell properties, *i*.*e*., long-term self-renewal, the ability to generate a large number of phenotypically distinct progenies, and tumor-initiating potential ^8^. Tumor cells with these features, defined as cancer stem cells, were subsequently identified in solid tumors, including GBM ^1-6^. The relevance of this paradigm is supported by the functional role of CSCs in tumor growth and recurrence and the association between stem cell signature (*i*.*e*., stemness) and poor patient prognosis ^9^. Diverse GBM cell populations defined phenotypically, based on the expression of markers such as CD133, CD44, ITGB8, PTPRZ1, or SOX2 ^7,9-16^, or based on fundamental functional characteristics, including being slow-cycling ^5,6,17^, exhibit the hallmark traits of CSCs. Although these cellular fractions may represent overlapping populations contributing to tumorigenesis, they can also define distinct lineages of cells or cellular states with different functions regulating tumor progression and treatment resistance. Are these lineages distinct from one another? Do they exhibit a unique profile of treatment resistance? The goal of this study is to address the question of CSC population heterogeneity with a specific focus on comparing slow-cycling cancer cells with cellular populations phenotypically characterized based on the expression of defined canonical CSC markers. We present phenotypic, genomic, and functional profiling, including comparing drug sensitivity.

## Material and Methods

### GBM patient-derived cell lines

Human GBM tumor specimens were cultured as previously described using the gliomasphere assay ^5,6,17,18^. Informed consent was acquired from patients and the experiments were conformed to and were approved by Institutional Review Board committee. Cells were cultured in serum-free conditions in NeuroCult NS-A Proliferation solution with 10% proliferation supplement (STEMCELL Technologies; Cat# 05750 and #05753) supplemented with 10 ng/ml basic fibroblast growth factor and 20 ng/ml human epidermal growth factor.

### Isolation of slow-cycling cells

Slow-cycling cells (SCCs) were isolated as described by Hoang and colleagues ^6^. Briefly, primary glioblastoma cells were labeled with CellTrace dye (Invitrogen) followed by a chase period of 5-10 days. Proliferation assessment of the cells was based on CellTrace fluorescence intensity decay rate over time measured by flow cytometry. SCCs were defined as the top 5-10% brightest cells.

### Antibodies

ITGB8 (R&D Systems, MAB4775), PTPRZ1 (BD Biosciences, 610179), SOX2 (R&D Systems, MAB2018), CD44 (Biolegend, 103007), CD133 (Miltenyi Biotec, 130-113-668).

### TMZ treatment

Cells were treated with 50uM or 500uM Temozolomide (TMZ, Sigma-Aldrich, T2577) 1 day after plating.

### Live/dead assay

After 3 days and 10 days of TMZ treatment, cocultured mCherry^+^/CD133^high^ cells and Wasabi^+^/SCCs were processed to single cells and labeled with fixable live/dead near-infrared fluorescent reactive dye (Invitrogen, L34975). The percentages of dead cells (live/dead dye^+^) were then compared through flow cytometry between CD133^high^ cells and SCCs in response to the different concentrations of TMZ.

### Lentivirus transduction

hGBM-L0 were transduced with vector pLV[exp]-CMV>mCherry (product ID LVS -VB191217-1841qsh-C, VectorBuilder) to constitutively express the fluorescent protein mCherry by following the manufacturer instructions. mCherry expressing transduced cells were isolated by flow cytometry.

### Flow cytometry

All flow cytometric studies were performed at the University of Florida Interdisciplinary Center for Biotechnology Flow Cytometry Core. The different cells populations were sorted out using a BD FACSAria II cytometer. BD LSR II or BD FACSymphony A3 cytometers were used for measuring and comparing cell viability and percentage of CD133^+^ ITGB8^+^, CD44^+^, PTPRZ1^+^, SOX2^+^ cells and SCCs.

### Bulk RNA sequencing

RNAseq was performed as previously described ^6^. SCCs isolated from nine different GBM patient-derived lines (L0, L1, L2, R24-01, R24-03, R24-23, R24-26, R24-37, R24-47) were sequenced for paired end 150 runs. Offline data were analyzed on the University of Florida High-Performance Cluster (HiPerGator). Briefly, low-quality reads and adaptors of fastq data were trimmed by trim_galore (Babraham Bioinformatics) then reads exceeding Q30 were aligned to Gencode v23 human genome by RSEM ^19^ to extract sample gene expression.

### Single-cell RNA sequencing

Single-cell RNA sequencing data were derived from Darmanis and colleagues ^20^. Malignant cells were selected (n=1091) based on the published metadata. Genes were considered positively expressed if the mean value of TMP > 0.2. Genes expressed in less than 30 cells were excluded. The group size was determined based on the expression distribution of the different CSC markers. The number of cells per group included in the study was defined by a homogenous range of max/min ratio (MMR) of expression lacking univariate outlier using box plot methods (**Supp. Table 1**). This identified the top 2% cells representing the CSC populations with the highest expression of each marker (*i*.*e*., 22 cells). Similarly, the top 2% cells for lipid metabolism and cell cycle score were used to define SCCs and FCCs, respectively. Escape package ^21^ was used for pathway enrichment analysis and the establishment of the lipid metabolism score. Cell cycle score was defined using Tirosh *et. al*. signature ^6,22,23^. G1S and G2M scores were defined and a new cell cycle score (CCS), based on the sum of G1S and G2M scores, was assigned to each cell. To visualize the level of cellular homology between groups, we used upset plots and Venn diagrams by Upset package ^24^ and Venny ^25^, respectively. Box plots for lipid metabolism signature score, CSC marker expression level, and cell cycle score were generated by ggplot2. Log10 or linear scale was applied based on the data distribution to achieve optimal visualization. Limma package ^26^ was used to identify differentially expressed genes (DEGs) between groups composed exclusively of private cells (n=22 for SCC, n=20 for FCC, n=19 for SOX2, n=18 for CD133, PTPRZ1, and CD44, and n=17 for ITGB8). p-values were adjusted using Bonferroni procedure, and the significance cutoff was set at 0.005. Ggplot2 and ggrepel packages ^27^ were used to generate volcano plots comparing gene expression levels between SCCs and each of the other groups. Uniform Manifold Approximation and Projection (UMAP) feature of Seurat 4.0 ^28,29^ was used as a deconvolution method to visualize the similarity or divergence between groups. Subsequently, trajectory analysis was integrated with Monocle 3 package. SCCs were set as a putative start point. A gradient color scale was applied to reflect pseudotime differences. The 1000 most variable genes were identified using CancerSubtypes package ^30^. Three-dimension principal component analysis was applied using Base -R. Hierarchical clustering was performed using pheatmap ^31^. The DEGs between SCC and CD133 were derived as described above. A drug target enrichment between SCC and CD133 was applied by Drugbank signature (version December 2021) ^32^ through Gene Set Enrichment Analysis, with a cutoff of False Discovery Rate (FDR) set at <0.05. Heatmap (heatmap.2 package) ^27^ was used to visualize the predicted drug sensitivity of the groups. All codes can be obtained upon request.

### Statistical tests

Wilcoxon rank sum test was applied for non-parametric pairwise comparison between reference group and each of the other groups (**Fig. 2**). One-way ANOVA combined with Bonferroni method were applied to compare cell death and cell ratio between different TMZ concentrations (**Fig. 4B-C, Supp. Fig. 4D**). p-values were adjusted for multiplicity using the Bonferroni method.

## Results

### Slow-cycling cells express a wide range of CSC markers

In GBM, we identified a subpopulation of cells displaying reduced cell cycle frequency and enriched in tumor-initiating and treatment-resistant cells exhibiting specific metabolism and enhanced infiltrative capacity ^6,33,34^. Demonstration of the stemness properties in SCCs and their progenies begs the question of how this lineage compares to the population of CSCs defined based on the expression of specific markers. Multiple experimental approaches can be used to identify, isolate, and study cancer SCCs (reviewed by Basu and colleagues) ^35^. We used label-retaining assays utilizing CellTrace dyes to interrogate these cells in GBM patient-derived lines ^6,17,33^ and evaluated by flow cytometry the expression of markers commonly used to enrich CSCs such as CD44, CD133, ITGB8, PTPRZ1, and SOX2. Our results indicate that although SCCs express markers of CSCs, not 100% of them are positive, revealing some phenotypic overlap and suggesting heterogeneity and distinction between these groups of cells (**Fig. 1A-B**). Additionally, we isolated SCCs by FACS from nine different primary GBM patient-derived cell lines and extracted RNA to be interrogated for bulk RNA sequencing analysis. In the SCCs, we observed a wide range of expression of the different CSC markers between the nine patients (**Fig. 1C**). Together, these data reveal that the property of being slow-cycling and the expression of canonical CSC markers do not seem to be mutually inclusive.

**Figure 1.**
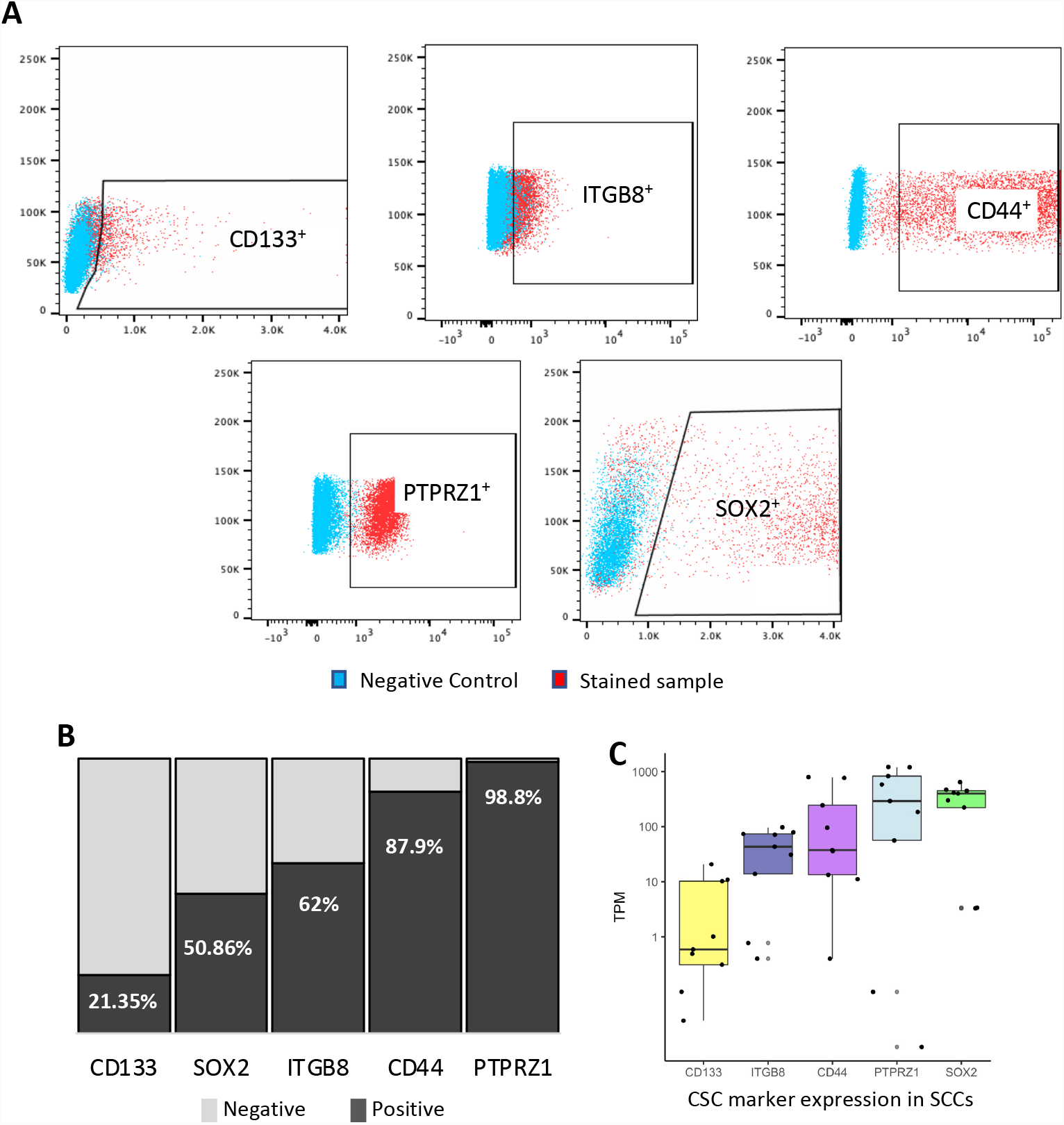
Expression level of CSCs markers in GBM SCCs. SCCs, identified as CellTrace retaining cells (top 5-10%) ^5,6^, were labeled with the following antibodies anti-CD133, ITGB8, CD44, PTPRZ1 and SOX2. Protein expression was measured by flow cytometry. **A)** Representative flow plots indicating the gates immunoreactive for the different CSC markers. **B)** Bar graph representing the percentage of SCCs that are positive (dark grey) or negative (light grey) for the different CSC markers. **C)** SCCs were FAC sorted from nine GBM patients and bulk RNA sequencing analysis was performed. The box plot indicates the level of CSC marker expression in SCCs for each patient, represented as transcript per million (TPM). The results identified SCCs in every patient and showed that SCCs exhibit a wide range of expression levels of CSC markers. Whiskers represent the 95% confidence interval and the box characterizes the interquartile range (IQR; 25^th^-50^th^-75^th^ percentiles).

### Single-cell transcriptomics identify multiple populations of CSCs

To further compare these populations of cells and quantify their state and potential dynamical structures we interrogated a single-cell RNA sequencing dataset ^20^ and defined SCCs based on the highest score (top 2%) of a recently reported metabolic signature (**Supp. Table 2**) ^6^, thereby identifying 22 cells (**Supp. Table 3**). Similarly, CSC populations were defined by the top 22 cells with the highest expression level of CD133, SOX2, PTPRZ1, ITGB8, or CD44 (**Supp. Table 3**). Fast-cycling cells (FCCs) were also included in our study and were delineated as the top 22 cells with the highest G1S/G2M cell cycle score, as previously described (**Supp. Table 3, Supp. Fig. 1A)** ^6,22,23^. We then compared the metabolic signature score, each CSC marker expression level, and cell cycle score between all populations as defined with the criteria described above. SCCs demonstrated a significantly greater lipid metabolism signature score than every other population (**Fig. 2A**). Interestingly, CD44^high^ cells exhibited the closest lipid score from SCCs compared to the classical CSC populations, with FCCs showing the furthest score from SCCs. Each CSC population (CD133^high^, SOX2^high^, PTPRZ1^high^, ITGB8^high^, CD44^high^) displayed significant overexpression of their respective marker compared to the other groups (**Fig. 2B-F**). Finally, FCCs demonstrated a higher cell cycle score than the other populations (**Fig. 2G**). Similar to the results from our bulk RNA sequencing studies presented on **figure 1C**, each of the 22 SCCs express heterogeneous expression levels of the CSC markers (CD133, SOX2, PTPRZ1, ITGB8, and CD44) ranging between several orders of magnitude (**Fig. 2H**).

**Figure 2.**
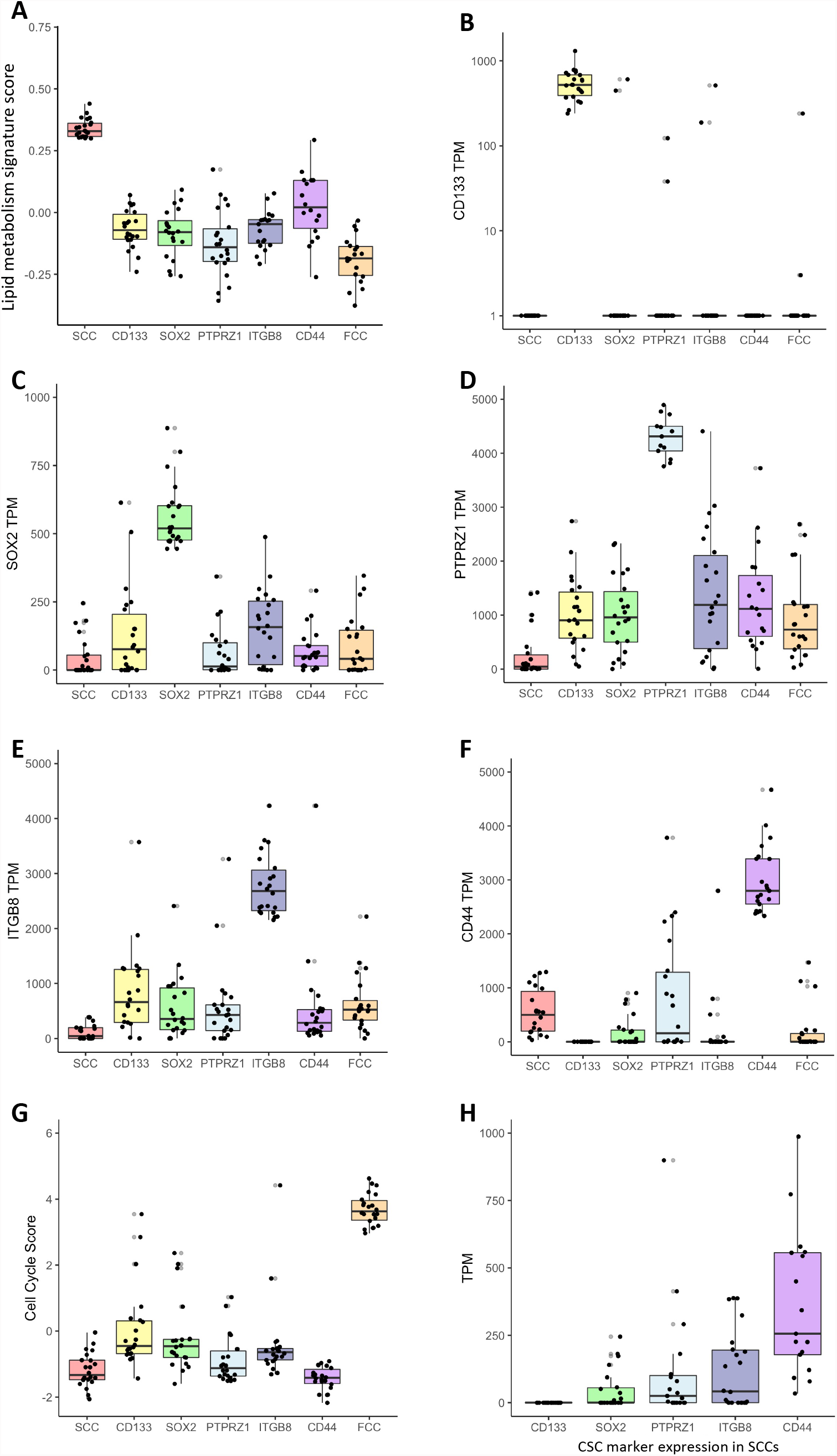
Gene signature scores and gene expression levels derived from scRNAseq comparing SCC, CSCs and FCC groups. Deconvolution score of lipid metabolism signature (**A**), expression of CD133 (**B**), SOX2 (**C**), PTPRZ1 (**D**), ITGB8 (**E**), CD44 (**F**), and cell cycle score (**G**). All pairwise comparisons comparing groups to the reference population (i.e., SCC-**A**, CD133-**B**, SOX2-**C**, PTPRZ1-**D**, ITGB8-**E**, CD44-**F**, FCC-**G**) were statistically significant (n=22, Wilcoxon test, all p-values adjusted for multiple comparisons using Bonferroni method were <0.001). error bars represent the 95% confidence interval and the box characterizes the IQR. **H)** Expression of CSC marker in SCCs.

### Heterogeneity between CSC populations

Each group was composed of 22 cells defining populations with limited (up to 7% between CD44^high^ and ITGB8^high^ groups) to no overlap (**Fig. 3A, Supp. Fig. 2**). Upset plot indicated that SCCs represent a unique population with none of the 22 cells being included in the other groups (CD133^high^, SOX2^high^, PTPRZ1^high^, ITGB8^high^, CD44^high^, and FCC), which all share at least one cell with each other (**Fig. 3A, Supp. Fig. 2**). Twenty cells were unique to FCCs, nineteen cells were exclusive to SOX2^high^ populations, whereas eighteen cells were specific to CD133^high^, PTPRZ1^high^, and CD44^high^ cells, and seventeen to ITGB8^high^ cells. Three cells were common between PTPRZ1^high^ and CD44^high^ groups. CD133^high^ and SOX2^high^ populations shared two cells. Finally, the following paired populations had one cell in common: CD133^high^/ITGB8^high^, SOX2^high^/ITGB8^high^, PTPRZ1^high^/ITGB8^high^, ITGB8^high^/CD44^high^, CD133^high^/FCC, and ITGB8^high^/FCC. We used the dimensionality reduction technique uniform manifold approximation and projection (UMAP) ^28,36^ for topological comparison of the cellular fractions. We found that both SCCs and CD44^high^ cells showed tight clustering, with CD44^high^ cells being the closest neighbors of SCCs, and CD133^high^ cells being the farthest ones (**Fig. 3B**). Even though cells were not harvested in a time series, they may be at different developmental stages. We therefore performed a trajectory analysis using Monocle3 ^37^ to model the potential relationships between groups of cells as a trajectory of gene expression changes. Interestingly, our pseudotemporal cell trajectory analysis placed SCCs at one end of pseudotime (close to CD44^high^ cells) and CD133^high^ cells at the opposite, divergent end. These results further support great distinctions between SCCs and CD133^high^ cells (**Fig. 3C**). Gene expression was compared between SCCs and the other cell populations. We identified sets of genes differentially regulated using the limma package ^26^ with a cutoff of log fold change (LogFC) greater than 2 or lower than −2. (**Supp. Table 4, Supp. Fig. 3**). A three-dimension principal component analysis was performed with the unique cells from each group using PCA scores calculated with the top 1000 variable genes across all populations. Three-dimensional imaging plotted to visualize the linear relationship between groups indicated that CD133^high^, SOX2^high^, PTPRZ1^high^, ITGB8^high^, and FCCs were closely distributed and distant from SCCs and CD44^high^ cells, which showed greater spreading and independent clustering (**Fig. 3D**). The expression level of these top 1000 variable genes was also represented as a heatmap, further illustrating the differential transcriptomic regulation between all of these populations (**Fig. 3E**).

**Figure 3.**
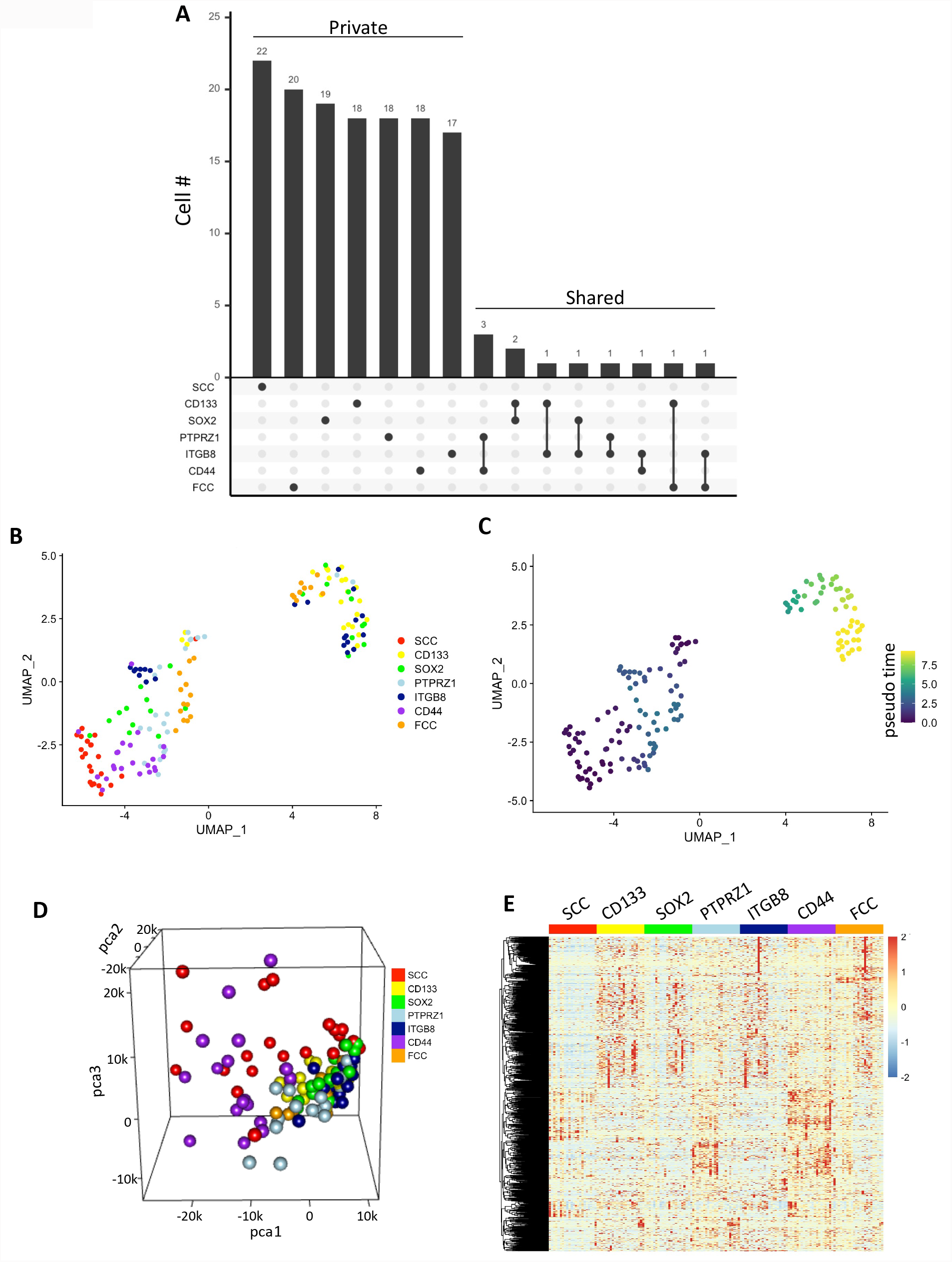
Transcriptomic differences between populations. **A)** Upset plot showing private or shared cells among groups. Set size is 22 cells for each group. **B)** UMAP projection of scRNA-seq data showing subsets of distinct cellular clusters. **C)** Trajectory analysis using Monocle3 coupled with Seurat single-cell data analysis package used for UMAP projection. **D)** Screenshot of a 3D-PCA using the top 1000 most variable genes. **E)** Heatmap displays groups’ hierarchical clustering using the top 1000 variable genes.

### Functional profiling of TMZ sensitivity

Temozolomide (TMZ) represents the standard-of-care chemotherapy used to treat GBM. The co-existence of phenotypically and functionally distinct subpopulations of cells exhibiting stemness properties may translate in remarkable heterogeneity of drug sensitivity. To address the question of specific drug response between cell populations, we functionally compared the effect of TMZ particularly between SCCs and CD133^high^ cells. These two cell populations were isolated from a primary GBM patient line (hGBM-L0) ^6,33^. mCherry-tagged CD133 cells and Wasabi-tagged SCCs were isolated by flow cytometry (**Fig. 4A, Supp. Fig.4A**) and co-cultured and treated with specific doses of TMZ (**Fig. 4B-D, Supp. Fig.4B-C**). Both populations of cells exhibit a different level of TMZ sensitivity, with CD133^high^ cells demonstrating significantly greater cell death compared to SCCs (**Fig. 4B, Supp. Fig.4B**), resulting in changes over time of the ratio SCC/CD133 in response to treatment (**Fig. 4C-D, Supp. Fig.4C-D**). These studies support the model of a heterogeneous pool of cells with CSC properties in GBM (*i*.*e*., SCCs vs. CD133^high^ cells), with a dynamic distribution that can be differentially regulated by therapies.

**Figure 4.**
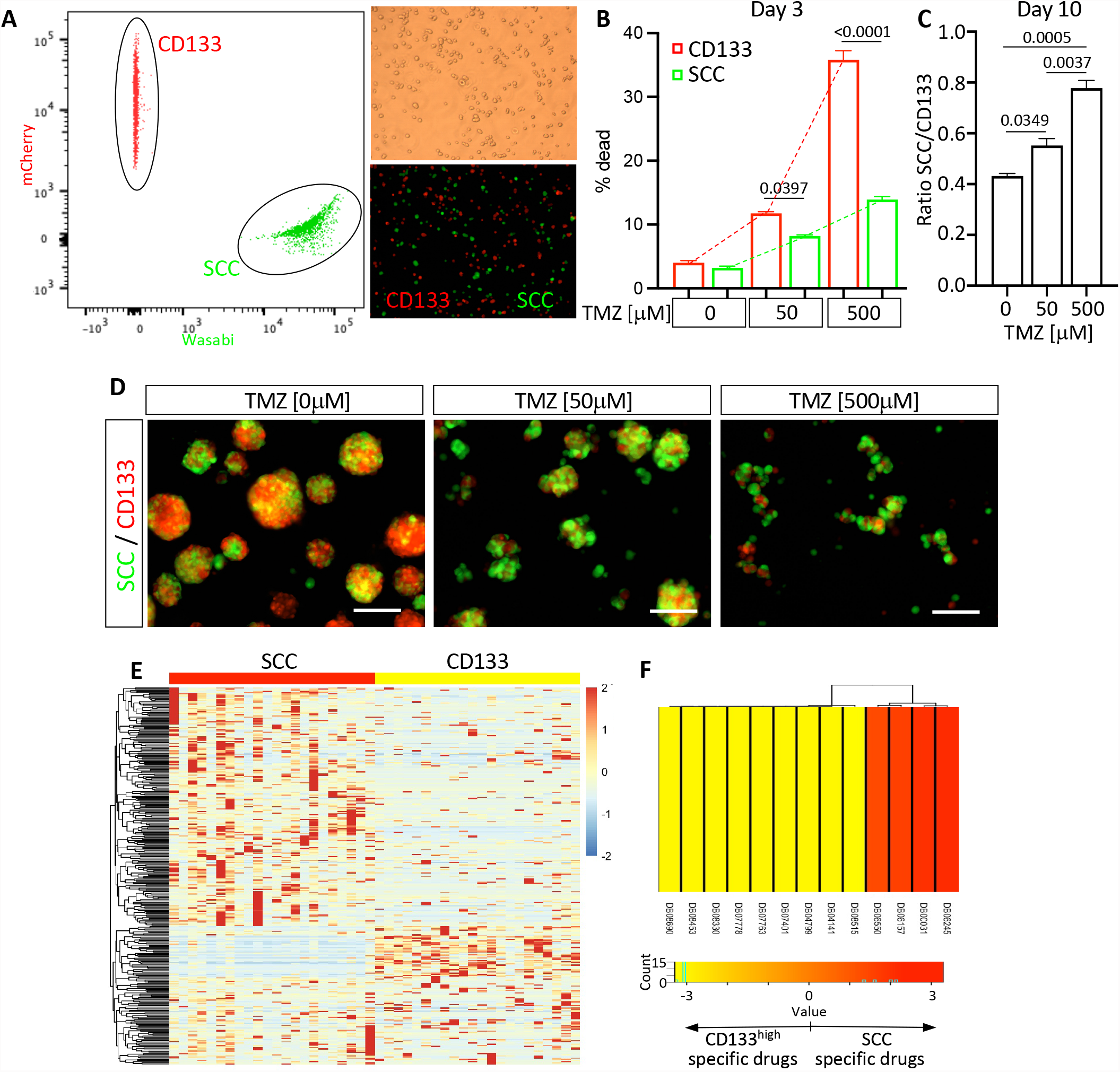
Functional assessment of the difference in drug sensitivity. **A)** Isolated from hGBM-L0, SCCs and CD133^high^ cells were co-cultured and treated with TMZ. **B)** Three days after initiating TMZ treatment, cell death was evaluated by flow cytometry using live/dead dye incorporation assay. Mean +/-SEM. one-way ANOVA. p-values were adjusted for multiplicity using the Bonferroni method. Results indicate distinct TMZ sensitivity between SCCs and CD133^high^ cells. **C)** Ten days after TMZ treatment, the ratio SCC/CD133 was compared between the experimental conditions. Results show a significant increase of the ratio, indicating a greater resistance to TMZ of SCCs compared to CD133^high^ cells. Mean +/-SEM. one-way ANOVA. p-values were adjusted for multiplicity using the Bonferroni method. **D)** Representative micrographs of co-cultured SCCs and CD133^+^ cells treated with TMZ. Scale bars, 100um. **E)** Hierarchical clustering using DEGs between SCCs and CD133^high^ cells. **F)** Drug target enrichment score between SCC and CD133^high^ groups. Drug IDs in red are agents specific to SCC. Drug IDs in yellow are specific to CD133^high^ cells. Drug ID: **DB06245**: Lanoteplase; **DB00031**: Tenecteplase; **DB06157**: Istaroxime; **DB06550**: Bivatuzumab; **DB08515**: (3AR,6R,6AS)-6-((S)-((S)-CYCLOHEX-2-ENYL) (HYDROXY)METHYL)-6A-METHYL-4-OXO-HEXAHYDRO-2H-FURO[3,2-C]PYRROLE-6-CARBALDEHYDE; **DB04141**: 2-Hexyloxy- 6-Hydroxymethyl-Tetrahydro-Pyran-3,4,5-Triol; **DB04799**: 6-Hydroxy-5-undecyl-4,7- benzothiazoledione; **DB07401**: Azoxystrobin; **DB07763**: (5S)-3-ANILINO-5-(2,4- DIFLUOROPHENYL)-5-METHYL-1,3-OXAZOLIDINE-2,4-DIONE; **DB07778**: (S)- famoxadone **DB08330**: METHYL (2Z)-3-METHOXY-2-{2-[(E)-2- PHENYLVINYL]PHENYL}ACRYLATE; **DB08453**: 2-Nonyl-4-quinolinol 1-oxide; **DB08690**:Ubiquinone Q2.

### Genomic profile predicting heterogeneity of drug-sensitivity

We used genomic profiling to further characterize the difference between SCCs and CD133^high^ cells and identify potential drugs predicted to target specifically SCCs versus CD133^high^ cells. DEGs between SCC and CD133^high^ groups (**Fig. 4E**) were used for drug target enrichment identification using the comprehensive online drugbank signature database through Gene Set Enrichment Analysis (GSEA) ^32^. Four drugs, including the humanized monoclonal antibody against CD44 Bivatuzumab, plasminogen activators Lanoteplase and Tenecteplase, as well as Na^+^/K^+^ ATPase inhibitor Istaroxime, were predicted to specifically target SCCs, whereas CD133^high^ cells were predicted to be sensitive to nine different other drugs (**Fig. 4F**). These results further illustrate the intratumoral functional differences between cell lineages and encourage us to functionally investigate the effect of these drugs in future studies.

## Discussion

Do populations of CSCs represent intermediate phenotypes along the spectrum of a single lineage? Do these cells present partial or complete functional redundancy with phenotypic distinction? Do CSCs exist as a homogeneous cellular population or do multiple CSCs co-exist in a given tumor? Do they reside at different stages or points of the same spectrum? Our study attempts to address these fundamental questions by further characterizing the CSC model in GBM.

Our results reveal differences in cell cycle kinetics, phenotypic and genomic profiles, and treatment sensitivity between multiple CSC populations and suggest distinct lineages or lineages with only partial overlap with a differential contribution to disease presentation and evolution. Specifically, our laboratory identified and characterized a subpopulation of slow-cycling cells in GBM ^6,17,33^. These cells represent a reservoir of tumor-initiating and treatment-resistant cells exhibiting CSC properties with the ability to give rise to highly proliferative progenies maintaining lineage specificity. Even though the progenies of SCCs can exhibit similar proliferative profiles to other cancer cell populations in response to specific cues and environment, their fate seems lineage-dependent and follow distinct transcriptional trajectories ^6,34^. The present study was designed to compare SCCs with established populations of CSCs. A combination of flow cytometry and bulk and single-cell RNA sequencing analyses revealed a substantial diversity of transcriptional profiles between SCCs and cells expressing the following CSC markers CD133, CD44, ITGB8, PTPRZ1 and SOX2. Together these results suggest that SCCs represent a distinct cell lineage with only a limited level of transcriptional redundancy. Of note, our study compared SCCs with only a few CSC populations; however additional markers could be selected, such as L1CAM, KLF4, integrin a6, ALDH, Nestin, Olig2, NANOG, ABCG2, or CD15 ^7,9,11,38-42^. Importantly, due to the lack of definite, universal and exclusive markers or functions identifying CSCs, discussion and controversy surrounding the conceptualization and contextualization of the cancer stem cell model and its hierarchical organization and regulation continue. We used a specific lipid metabolism signature, which we previously demonstrated to identify slow-cycling cells and their progenies, to classify cells as SCCs from a single cell RNA sequencing dataset ^20^. We compared these cells with different CSCs populations defined based on the high expression of canonical CSC markers (*i*.*e*., CD133, ITGB8, CD44, PTPRZ1, and SOX2). Our results showed a lack of cellular overlap between SCCs and the other CSC populations, further supporting lineage specificity (**Fig. 3A, Supp. Fig. 2**).

Using a coculture dual-color system, in which CD133^+^ cells were tagged with the fluorescent protein mCherry and SCCs with the fluorescent protein Wasabi, we were able to compare in real time lineage dynamics in response to treatment. Our results indicated that SCCs and their progenies are more tolerant to TMZ than the CD133^high^ cell lineage (**Fig. 4D-F, Supp. Fig. 4B-D**). Similarly, Reinartz and colleagues demonstrated specific subclone dynamism and functional consequences of intratumoral heterogeneity of drug resistance in GBM ^43^. This work also supports GBM as a disease with the co-existence of polyclonal collections of cellular hierarchies combining cancer stem cell and classical stochastic models. Oren and colleagues used a high-complexity expressed barcode lentiviral library for simultaneous tracing of cell clonal origin and proliferative and transcriptional profiling. Their results show the existence of treatment-resistant persisters in lung cancers, with their fate being lineage-dependent and characterized by metabolic reprogramming of anti-oxidant and lipid pathways ^44^. This data is in line with our previous study demonstrating that, under treatment pressure, slow-cycling cancer stem cells give rise to lineage-specific cycling persisters that repopulate the tumors and are also marked by up-regulated fatty acid metabolic pathways and anti-oxidant programs ^6^. These pathways, especially autophagy and lipid droplets metabolism that we reported being increased in SCCs ^6^, represent candidate regulators of diapause ^45-47^, which is a potential mechanism by which SCCs may enter or exhibit drug-tolerant persister (DTP) state ^48,49^. Diapause is a defined state of physiological dormancy characterized by a dormant stage of suspended embryonic development triggered by stress. Two recent studies suggest that tumor cells can engage diapause-like pathways enabling cancer treatment escape ^48,49^. Rehman and colleagues reported that colorectal cancer cells are equipotent in their ability to enter the DTP state by activating diapause-like transcriptional programs to survive therapy. Conversely, our data suggest a different scenario in GBM, which display great heterogeneity characterized by diverse populations of cells, especially cancer stem cells with distinct treatment sensitivity, suggesting a varied capacity to stimulate diapause-like mechanisms to enter the DTP state. Considering this heterogeneity, therapeutic strategies aiming to eliminate cells with stemness properties will have to be combinatorial and target every individual lineage.

Diversity in CSCs populations is now well recognized. However, the precise hierarchical organization and the plasticity of this organization between CSC populations and non-CSCs are very complex and challenging to appreciate and understand fully. The potential hierarchical link between SCCs and the classical CSCs could be further investigated using lineage tracing assays, similar to the report by Lan and colleagues ^50^. This study used DNA barcoding and fate mapping to demonstrate a model with functionally distinct cells in GBM with a conserved proliferative hierarchy in which slow-cycling stem-like cells give rise to rapidly cycling progenitors, showing extensive self-renewing capability with the ability to generate terminally differentiated cells.

GBM are spatially organized complex ecosystems with heterogeneity across the tumor microenvironment, where specific CSC lineages may be selected based on their spatial distribution within the tumor ^9^, defining niche-specific cell-cell interactions. Understanding of the dynamic transcriptional and spatial fluctuations of each CSC lineage, and the interconnection and interconversion of these populations will be paramount for developing precision and effective therapies. The use of sophisticated high-throughput approaches, which may combine mathematical modeling, artificial intelligence, single-cell RNA sequencing, 3D model systems, multiplex imaging, and spatial transcriptomics, will help map and understand this dynamically adaptive complex system and uncover the mechanisms underlying its resilience that is the root of its resistance to treatment.

## Abbreviations

(SCC): slow-cycling cell
(FCC): fast-cycling cell
(CSC): cancer stem cell
(CCS): cell cycle score
(TPM): transcript per million
(DTP): drug-tolerant persister

## Acknowledgements

The authors would like to thank Dr. Chang’s laboratory at the University of Florida for kindly providing hGBM-L0 mWasabi expressing cells. Funding for this research was supported in part by grants from the NIH (R21NS116578 to L.P.D., R24NS086554 to B.A.R.), the Florida Center for Brain Tumor Research and Accelerate Brain Cancer Cure (to L.P.D.), the University of Florida Health Cancer Center (Pilot Grant to L.P.D.), and the St. Baldrick’s Foundation (to L.P.D.).

## Supplemental Figure/Table Legends

**Supplemental Figure 1.**
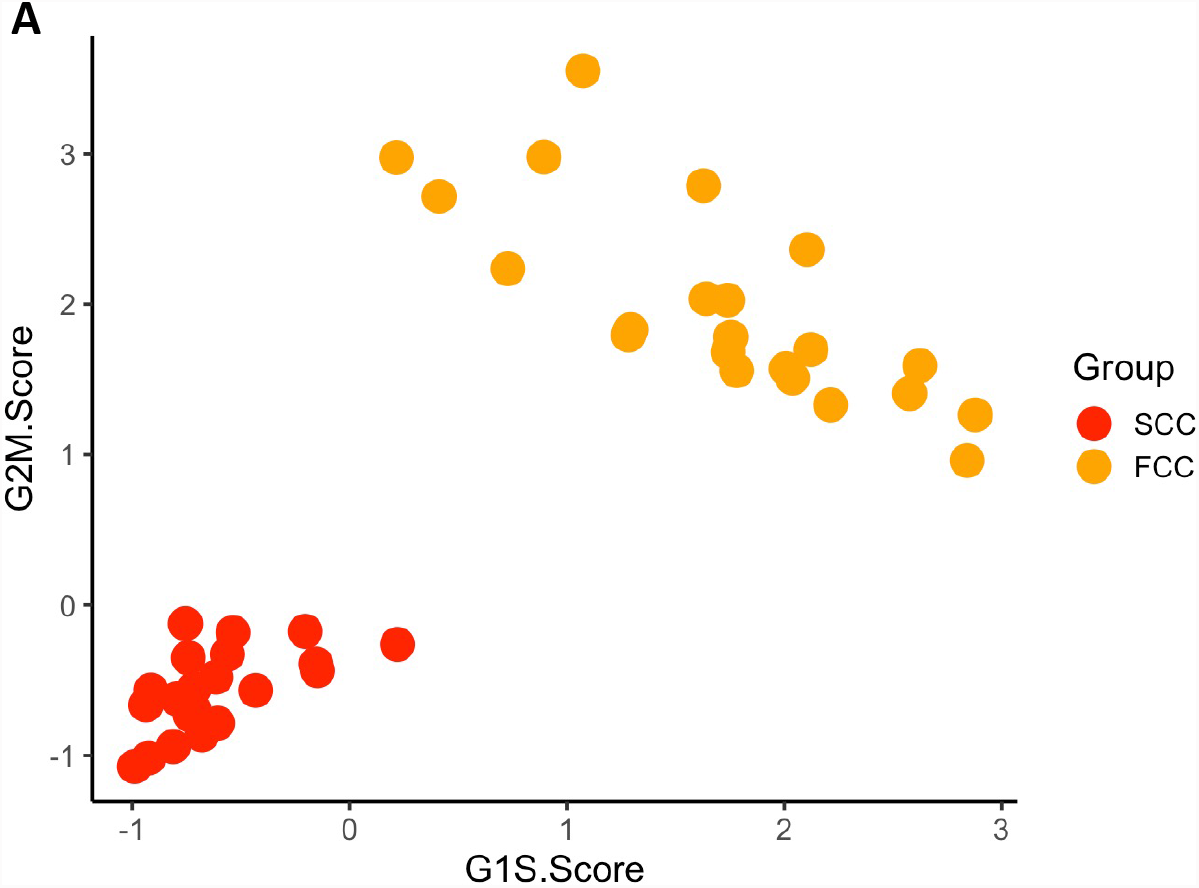
(related to Figure 2). Cell cycle score. Scatter plot representing the cell cycle scores of SCCs (red, n=22) and FCCs (orange, n=22).

**Supplemental Figure 2.**
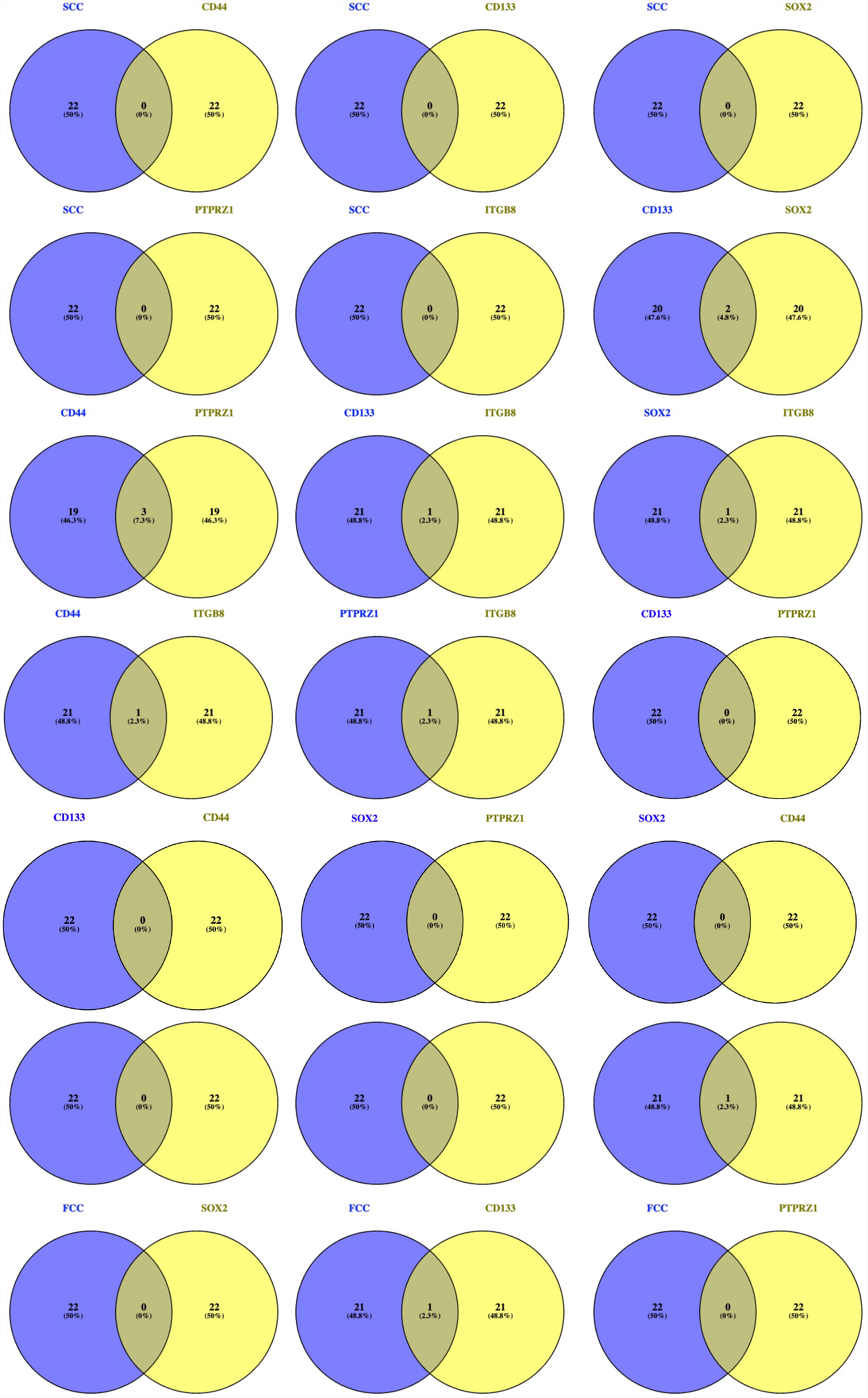
(related to Figure 3). Pairwise Venn Diagrams. The numbers in the overlapped areas of each Venn Diagram represent the number of cells in common between the two groups. The non-overlapping sections show the number of cells private to each population.

**Supplemental Figure 3.**
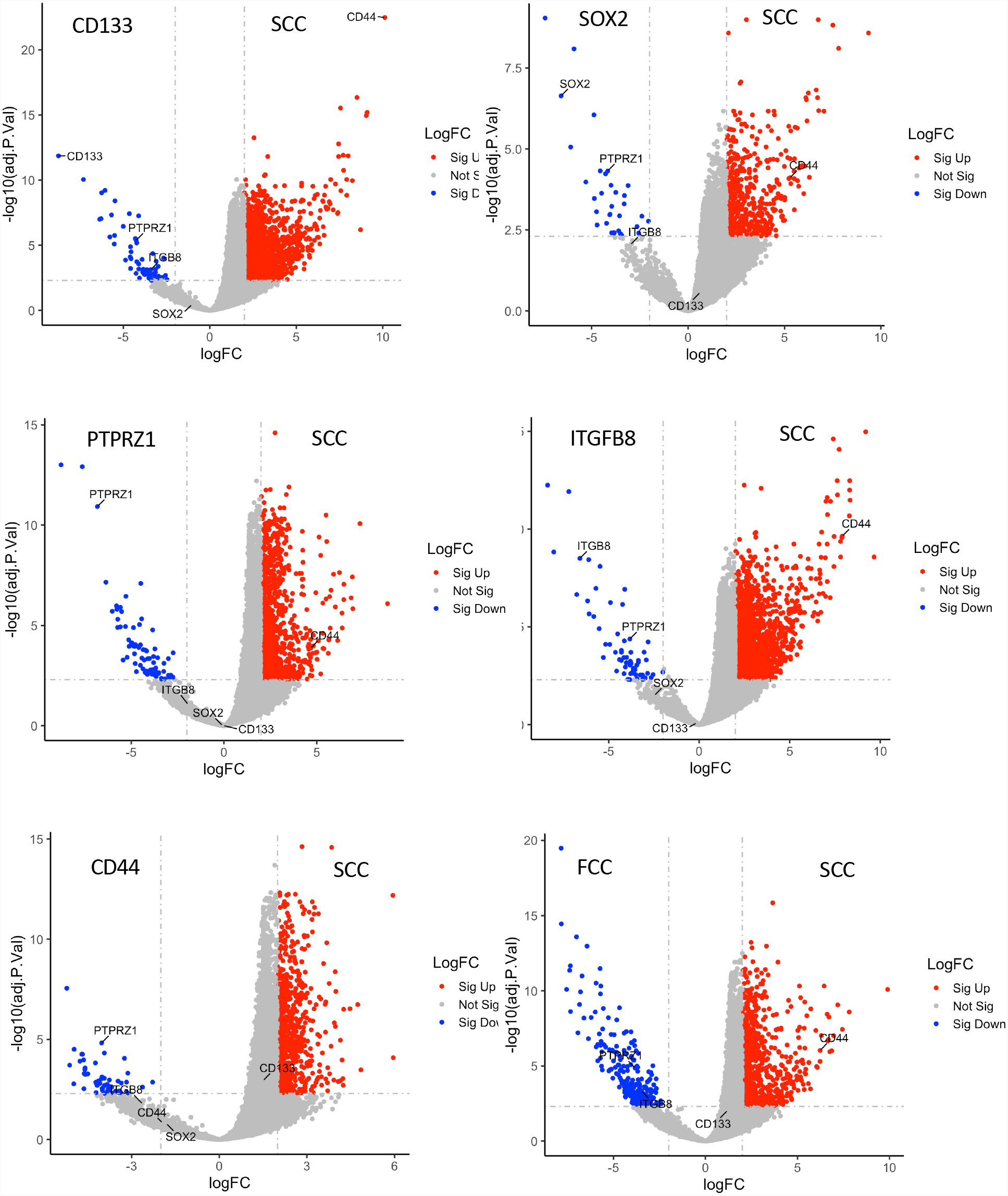
(related to Figure 3). Volcano plots comparing gene expression between SCC and each of the other groups. Red dots represent significantly upregulated genes in SCCs. Blue dots are significantly downregulated genes in SCCs. CSC markers (CD133, SOX2, PTPRZ1, ITGB8, CD44) were annotated. Differentially Expressed Genes (DEGs) were filtered based on LogFC over than 2 or less than −2 with an adjusted p-value set as 0.05.

**Supplemental Figure 4.**
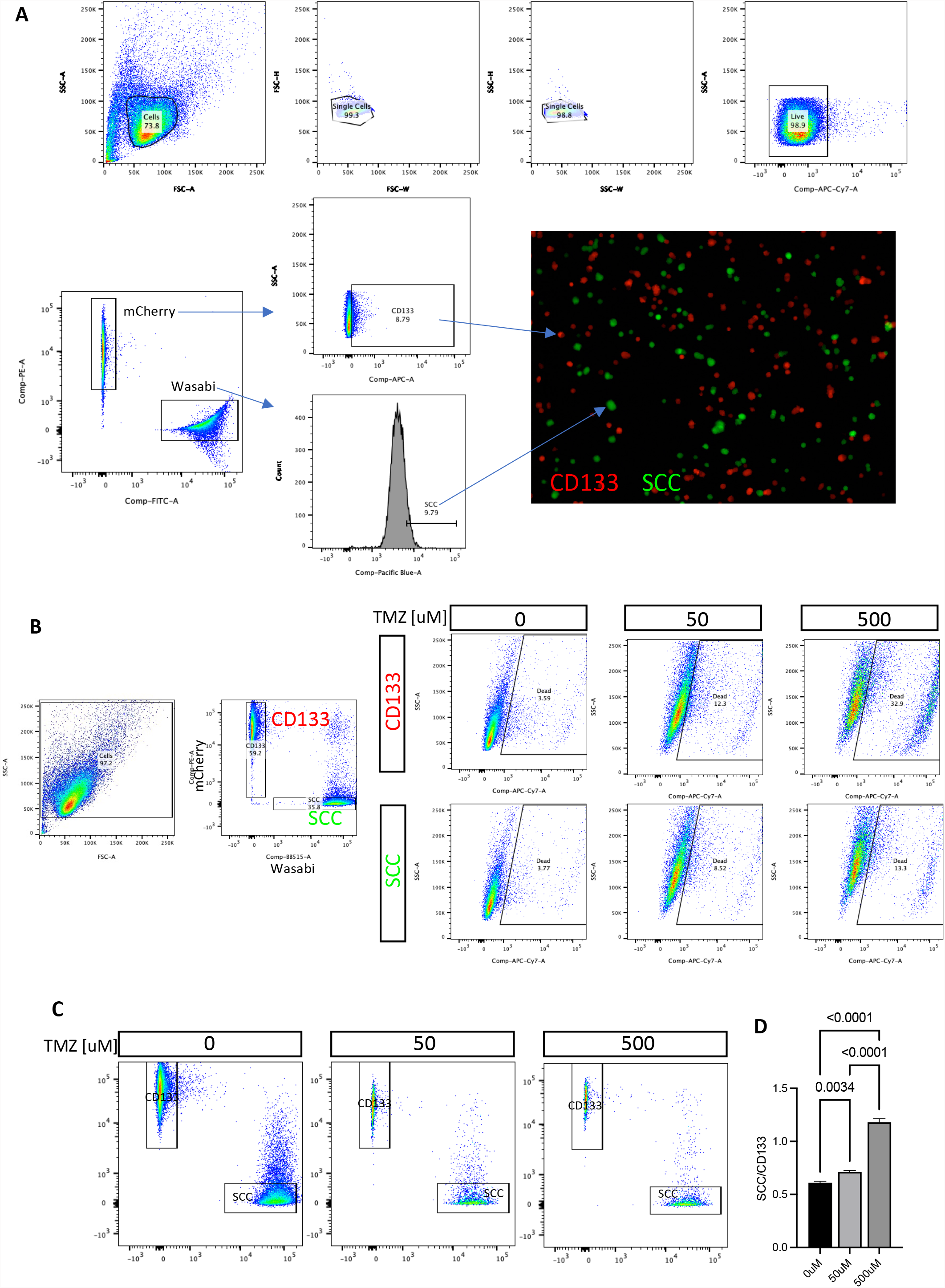
(related to Figure 4). Gating strategy for sorting CD133^+^ cells and SCCs and functional TMZ sensitivity assay. **A)** Primary hGBM-L0 cells ^5,6^ were transduced to constitutively express the fluorescent reporter tag Wasabi or mCherry. Wasabi expressing cells were labeled with CellTrace Violet and chased for one week before isolating SCCs (top 10%). mCherry-tagged CD133 immunoreactive cells were also FAC sorted and co-cultured with wasabi-tagged SCCs at an initial 40/60 ratio of SCC/CD133. The cells were then cultured and treated with various doses of TMZ. **B)** Gating strategy to measure cell death. The percentage of dead cells was quantified by flow cytometry after incubating the cultures with live/dead fixable reactive dye. The gating strategy is presented. **C)** The ratio of SCC/CD133 cells was evaluated by quantifying using flow cytometry the percentage of wasabi^+^ cells and mCherry^+^ cells in the different experimental groups. Representative flow dot plots are presented. **D)** Graph represents the mean of the ratio wasabi/mCherry, measured three days post-TMZ treatment, +/-SEM, one-way ANOVA. p-values were adjusted for multiplicity using the Bonferroni method.

**Supplemental Table 1**. List of max/min ratio (MMR) of expression of each CSC marker.

**Supplemental Table 2**. List of 56 genes defining SCC-enriched lipid metabolism signature ^6^.

**Supplemental Table 3**. Complete list of malignant cell ID identified from Darmanis and colleagues ^20^ with the level of expression of CD133, ITGB8, CD44, PTPRZ1, and SOX2, as well as lipid metabolism signature and cell cycle scores.

**Supplemental Table 4**. List of genes differentially regulated between SCCs and the other cell populations extracted from scRNA Seq dataset ^20^.

## References

1 Ignatova, T. N. et al. Human cortical glial tumors contain neural stem-like cells expressing astroglial and neuronal markers in vitro. Glia 39, 193–206 (2002).

2 Galli, R. et al. Isolation and characterization of tumorigenic, stem-like neural precursors from human glioblastoma. Cancer Res 64, 7011–7021 (2004).

3 Singh, S. K. et al. Identification of human brain tumour initiating cells. Nature 432, 396–401 (2004).

4 Bao, S. et al. Glioma stem cells promote radioresistance by preferential activation of the DNA damage response. Nature 444, 756–760, doi:10.1038/nature05236 (2006).

5 Deleyrolle, L. P. et al. Evidence for label-retaining tumour-initiating cells in human glioblastoma. Brain : a journal of neurology 134, 1331–1343, doi:10.1093/brain/awr081 (2011).

6 Hoang-Minh, L. B. et al. Infiltrative and drug-resistant slow-cycling cells support metabolic heterogeneity in glioblastoma. EMBO J 37, doi:10.15252/embj.201798772 (2018).

7 Mitchell, K., Troike, K., Silver, D. J. & Lathia, J. D. The evolution of the cancer stem cell state in glioblastoma: emerging insights into the next generation of functional interactions. Neuro Oncol 23, 199–213, doi:10.1093/neuonc/noaa259 (2021).

8 Lapidot, T. et al. A cell initiating human acute myeloid leukaemia after transplantation into SCID mice. Nature 367, 645–648, doi:10.1038/367645a0 (1994).

9 Prager, B. C., Bhargava, S., Mahadev, V., Hubert, C. G. & Rich, J. N. Glioblastoma Stem Cells: Driving Resilience through Chaos. Trends Cancer 6, 223–235, doi:10.1016/j.trecan.2020.01.009 (2020).

10 Bhaduri, A. et al. Outer Radial Glia-like Cancer Stem Cells Contribute to Heterogeneity of Glioblastoma. Cell stem cell 26, 48–63 e46, doi:10.1016/j.stem.2019.11.015 (2020).

11 Lathia, J. D., Mack, S. C., Mulkearns-Hubert, E. E., Valentim, C. L. & Rich, J. N. Cancer stem cells in glioblastoma. Genes & development 29, 1203–1217, doi:10.1101/gad.261982.115 (2015).

12 Gangemi, R. M. et al. SOX2 silencing in glioblastoma tumor-initiating cells causes stop of proliferation and loss of tumorigenicity. Stem cells 27, 40–48, doi:10.1634/stemcells.2008-0493 (2009).

13 Berezovsky, A. D. et al. Sox2 promotes malignancy in glioblastoma by regulating plasticity and astrocytic differentiation. Neoplasia 16, 193-206, 206 e119-125, doi:10.1016/j.neo.2014.03.006 (2014).

14 Shi, Y. et al. Tumour-associated macrophages secrete pleiotrophin to promote PTPRZ1 signalling in glioblastoma stem cells for tumour growth. Nature communications 8, 15080, doi:10.1038/ncomms15080 (2017).

15 Xia, Z. et al. The Expression, Functions, Interactions and Prognostic Values of PTPRZ1: A Review and Bioinformatic Analysis. J Cancer 10, 1663–1674, doi:10.7150/jca.28231 (2019).

16 Guerrero, P. A. et al. Glioblastoma stem cells exploit the alphavbeta8 integrin-TGFbeta1 signaling axis to drive tumor initiation and progression. Oncogene 36, 6568–6580, doi:10.1038/onc.2017.248 (2017).

17 Deleyrolle, L. P., Rohaus, M. R., Fortin, J. M., Reynolds, B. A. & Azari, H. Identification and isolation of slow-dividing cells in human glioblastoma using carboxy fluorescein succinimidyl ester (CFSE). J Vis Exp, doi:10.3791/3918 (2012).

18 Sarkisian, M. R. et al. Detection of primary cilia in human glioblastoma. Journal of neuro-oncology 117, 15–24, doi:10.1007/s11060-013-1340-y (2014).

19 Li, B. & Dewey, C. N. RSEM: accurate transcript quantification from RNA-Seq data with or without a reference genome. BMC Bioinformatics 12, 323, doi:10.1186/1471-2105-12-323 (2011).

20 Darmanis, S. et al. Single-Cell RNA-Seq Analysis of Infiltrating Neoplastic Cells at the Migrating Front of Human Glioblastoma. Cell reports 21, 1399–1410, doi:10.1016/j.celrep.2017.10.030 (2017).

21 Borcherding, N. & Andrews, J. (2020).

22 Tirosh, I. et al. Dissecting the multicellular ecosystem of metastatic melanoma by single-cell RNA-seq. Science 352, 189–196, doi:10.1126/science.aad0501 (2016).

23 Tirosh, I. et al. Single-cell RNA-seq supports a developmental hierarchy in human oligodendroglioma. Nature 539, 309-+, doi:10.1038/nature20123 (2016).

24 Conway, J. R., Lex, A. & Gehlenborg, N. UpSetR: an R package for the visualization of intersecting sets and their properties. Bioinformatics 33, 2938–2940, doi:10.1093/bioinformatics/btx364 (2017).

25 Oliveros, J. C. Venny. An interactive tool for comparing lists with Venn’s diagrams. (2007).

26 Ritchie, M. E. et al. limma powers differential expression analyses for RNA-sequencing and microarray studies. Nucleic acids research 43, e47, doi:10.1093/nar/gkv007 (2015).

27 Wickham, H. ggplot2: Elegant Graphics for Data Analysis. (2016).

28 McInnes, L. & Healy, J. UMAP: uniform manifold approximation and projection for dimension reduction.. Preprint at https://arxiv.org/abs/1802.03426 (2018).

29 Hao, Y. et al. Integrated analysis of multimodal single-cell data. Cell 184, 3573-3587.e3529, doi:10.1016/j.cell.2021.04.048 (2021).

30 Xu, T. et al. CancerSubtypes: an R/Bioconductor package for molecular cancer subtype identification, validation and visualization. Bioinformatics 33, 3131–3133, doi:10.1093/bioinformatics/btx378 (2017).

31 Kolde, R. (2015).

32 Merico, D., Isserlin, R., Stueker, O., Emili, A. & Bader, G. D. Enrichment map: a network-based method for gene-set enrichment visualization and interpretation. PLoS One 5, e13984, doi:10.1371/journal.pone.0013984 (2010).

33 Deleyrolle, L. P. et al. Evidence for label-retaining tumour-initiating cells in human glioblastoma. Brain 134, 1331–1343, doi:10.1093/brain/awr081 (2011).

34 Badr, C. E., Silver, D. J., Siebzehnrubl, F. A. & Deleyrolle, L. P. Metabolic heterogeneity and adaptability in brain tumors. Cell Mol Life Sci 77, 5101–5119, doi:10.1007/s00018-020-03569-w (2020).

35 Basu, S., Dong, Y., Kumar, R., Jeter, C. & Tang, D. G. Slow-cycling (dormant) cancer cells in therapy resistance, cancer relapse and metastasis. Semin Cancer Biol, doi:10.1016/j.semcancer.2021.04.021 (2021).

36 Becht, E. et al. Dimensionality reduction for visualizing single-cell data using UMAP. Nat Biotechnol, doi:10.1038/nbt.4314 (2018).

37 Trapnell, C. et al. The dynamics and regulators of cell fate decisions are revealed by pseudotemporal ordering of single cells. Nat Biotechnol 32, 381–386, doi:10.1038/nbt.2859 (2014).

38 Rasper, M. et al. Aldehyde dehydrogenase 1 positive glioblastoma cells show brain tumor stem cell capacity. Neuro Oncol 12, 1024–1033, doi:10.1093/neuonc/noq070 (2010).

39 Jin, X., Jin, X., Jung, J. E., Beck, S. & Kim, H. Cell surface Nestin is a biomarker for glioma stem cells. Biochem Biophys Res Commun 433, 496–501, doi:10.1016/j.bbrc.2013.03.021 (2013).

40 Ligon, K. L. et al. Olig2-regulated lineage-restricted pathway controls replication competence in neural stem cells and malignant glioma. Neuron 53, 503–517, doi:10.1016/j.neuron.2007.01.009 (2007).

41 Zbinden, M. et al. NANOG regulates glioma stem cells and is essential in vivo acting in a cross-functional network with GLI1 and p53. EMBO J 29, 2659–2674, doi:10.1038/emboj.2010.137 (2010).

42 Son, M. J., Woolard, K., Nam, D. H., Lee, J. & Fine, H. A. SSEA-1 is an enrichment marker for tumor-initiating cells in human glioblastoma. Cell stem cell 4, 440–452, doi:10.1016/j.stem.2009.03.003 (2009).

43 Reinartz, R. et al. Functional Subclone Profiling for Prediction of Treatment-Induced Intratumor Population Shifts and Discovery of Rational Drug Combinations in Human Glioblastoma. Clinical cancer research : an official journal of the American Association for Cancer Research 23, 562–574, doi:10.1158/1078-0432.CCR-15-2089 (2017).

44 Oren, Y. et al. Cycling cancer persister cells arise from lineages with distinct programs. Nature 596, 576–582, doi:10.1038/s41586-021-03796-6 (2021).

45 Hussein, A. M. et al. Metabolic Control over mTOR-Dependent Diapause-like State. Dev Cell 52, 236–250 e237, doi:10.1016/j.devcel.2019.12.018 (2020).

46 Tan, Q. Q., Liu, W., Zhu, F., Lei, C. L. & Wang, X. P. Fatty acid synthase 2 contributes to diapause preparation in a beetle by regulating lipid accumulation and stress tolerance genes expression. Scientific reports 7, 40509, doi:10.1038/srep40509 (2017).

47 Arena, R. et al. Lipid droplets in mammalian eggs are utilized during embryonic diapause. Proceedings of the National Academy of Sciences of the United States of America 118, doi:10.1073/pnas.2018362118 (2021).

48 Dhimolea, E. et al. An Embryonic Diapause-like Adaptation with Suppressed Myc Activity Enables Tumor Treatment Persistence. Cancer Cell 39, 240–256 e211, doi:10.1016/j.ccell.2020.12.002 (2021).

49 Rehman, S. K. et al. Colorectal Cancer Cells Enter a Diapause-like DTP State to Survive Chemotherapy. Cell 184, 226–242 e221, doi:10.1016/j.cell.2020.11.018 (2021).

50 Lan, X. Y. et al. Fate mapping of human glioblastoma reveals an invariant stem cell hierarchy. Nature 549, 227-+, doi:10.1038/nature23666 (2017).

